# pLM-SAV: A Δ-Embedding Approach for Predicting Pathogenic Single Amino Acid Variants

**DOI:** 10.1101/2025.05.24.655916

**Authors:** Orsolya Gereben, Hedvig Tordai, Lana Khamisi, Erda Qorri, Tamás Hegedűs

## Abstract

Predicting whether single amino acid variants (SAVs) in proteins lead to pathogenic outcomes is a critical challenge in molecular biology and precision medicine. Experimental determination of all possible mutation effects is infeasible, and while state-of-the-art tools such as AlphaMissense show promise, their diagnostic performance is insufficient and they are often difficult to run locally. We developed pLM-SAV, a simple yet effective predictor that leverages protein language models (pLMs). Δ-embeddings, computed as the difference between wild-type and mutant sequence embeddings, are used as input for a convolutional neural network. To prevent data leakage, we trained our model on a well-characterized, labeled set of Eff10k and evaluated it on a non-homologous subset of ClinVar data. Our results demonstrate that this approach performs exceptionally well on the Eff10k test folds and reasonably on ClinVar test sets. Notably, pLM-SAV excels in resolving ambiguous predictions by AlphaMissense. We also found that an ensemble method, REVEL, outperforms both AlphaMissense and pLM-SAV, thus, we integrated these REVEL- enhanced predictions into our widely used AlphaMissense web application. Our results demonstrate that an SAV predictor trained on labeled data can achieve high predictive performance. Unlike previous methods such as VESPA, pLM-SAV uses no handcrafted features or substitution matrices, relying solely on pLM-derived representations. We anticipate that incorporating delta-embeddings into other mutation effect predictors or mutant structure prediction methods will further enhance their accuracy and utility in diverse biological contexts.

**Availability and Implementation:** Freely available at https://doi.org/10.5281/zenodo.15502498 and https://alphamissense.hegelab.org.

## Introduction

Understanding the functional consequences of single amino acid variants (SAVs) is a major goal in molecular biology, human genetics, and precision medicine [1,2]. SAVs can alter protein structure, dynamics, interactions, or stability, and are often associated with human disease [3,4]. As high-throughput sequencing continues to uncover vast numbers of novel variants, which include many variants of uncertain significance (VUS), reliable in silico tools are needed to predict whether a given amino acid substitution is likely to be pathogenic or benign. However, experimental validation of all possible variants is impractical, and often impossible, especially for large protein families or non-model organisms [5,6].

Over the past decade, many computational methods have been developed to estimate the functional impact of SAVs. Among the most widely used are rule-based and supervised learning models that combine evolutionary conservation with structural, biophysical, or annotation-derived features. An important early method is SNAP2 [7]. SNAP2 is a supervised predictor trained using 10-fold cross-validation on the Eff10k dataset. It combines a wide range of features, including evolutionary and structural data, such as sequence alignments, secondary structure, solvent accessibility, and biophysical properties. It also integrates curated annotations and functional signals from resources like SWISS-Prot, Pfam, PROSITE, and AAindex, making it a feature-rich and biologically informed model.

REVEL (Rare Exome Variant Ensemble Learner) [8] is a powerful ensemble predictor combining multiple tools to prioritize likely pathogenic variants, and has been widely adopted in clinical genomics pipelines. DeepMind’s AlphaMissense [9] provides predictions for 71 million single-nucleotide missense variants and a superset of 216 million possible amino acid substitutions. It builds on the AlphaFold2 framework, using structure-based features to train a deep learning model. AlphaMissense has shown strong performance on standard benchmarks such as ClinVar and ProteinGym, but its diagnostic utility is still limited. Moreover, it is restricted to human proteins, and its models are not easily deployed or retrained locally.

Among the more recent generation of embedding-based predictors is VESPA, developed by Marquet et al. [10]. VESPA infers conservation using protein language model (pLM) embeddings and supplements this information with BLOSUM substitution scores and amino acid substitution probabilities. Importantly, VESPA and its variant VESPAl were trained on the Eff10k dataset using a methodology similar to ours in this study, making them highly relevant for direct comparison. Several methods have been developed that exploit pLMs in diverse ways to predict SAV effects [11,12].

The emergence of protein language models (pLMs) has transformed computational biology by enabling high-quality sequence embeddings learned from massive databases of unlabeled protein sequences. These models, such as ProtT5 [13] and Ankh [14] capture rich contextual and evolutionary information without the need for multiple sequence alignments. Protein embeddings generated by pLMs have already shown promise for downstream tasks, including structure prediction, function annotation, and variant effect estimation [15,16].

Here, we explore whether delta-embeddings — the difference between the embedding vectors of the wild-type and mutant protein sequences — encode meaningful information for SAV effect prediction. While embeddings from pLMs implicitly capture residue-level context, it is unclear whether simple subtraction can expose mutation-specific effects that are useful for classification. To address this, we developed pLM-SAV, a lightweight convolutional neural network (CNN) model trained on delta-embeddings derived from either ProtT5 or Ankh [13,14]. The model was trained on a clustered, non-redundant subset of the Eff10k dataset [7,10] and evaluated on ClinVar proteins with no sequence similarity to the training data, ensuring robust generalization. Additionally, we assessed pLM-SAV’s performance on ambiguous cases predicted by AlphaMissense and REVEL, and compared it to other predictors on the ProteinGym benchmark.

## Methods

### Datasets

Eff10k, a dataset originally developed in the Rost laboratory (https://www.rostlab.org) for the SNAP2 and VESPA predictors [7,10], was obtained from the authors along with their predefined fold split. The set was combined from three sources: PMD (the Protein Mutant Database [17], Protein specific studies [18,19], and disease variants form OMIM (Online Mendelian Inheritance in Men) [20,21], as detailed in [7]. The authors clustered protein sequences using PSI-BLAST with an E-value threshold of <1e–3 to group homologous proteins. The cluster sizes greatly varied (1 to 1,941 members) [7]. These clusters were then randomly assigned to 10 folds, resulting in one disproportionately large fold (Fold 0, containing 44,024 variants) and nine approximately equal-sized folds. This clustering and folding strategy ensured that no homologous sequences were shared across folds, thereby preserving the integrity of cross-validation and preventing data leakage. We adopted this fold assignment unchanged for comparability with SNAP2 and VESPA. The full dataset contains 100,737 binary SAV-effect annotations from 9,594 proteins, of which 99,926 were used in our analysis (39,299 classified as neutral and 60,627 labeled as effect) with an effect-to-neutral ratio of approximately 1.5.

PMD4k is a subset of the Eff10k dataset derived from the Protein Mutant Database (PMD) [17]. It has been used for testing in previous SAV effect prediction methods, including SNAP2 [7] and VESPA [10]. The dataset comprises 51,266 variants, including 13,497 classified as neutral and 37,769 as effect. It can be further divided into 676 human proteins with 2,788 neutral and 6,863 effect variants, and 3,343 non-human proteins with 10,709 neutral and 30,906 effect variants. To prevent data leakage, we followed the approach of Rost and colleagues [7,10] and evaluated predictions only for PMD4k variants in the test fold of each model. Since PMD4k is a subset of Eff10k, the PMD variants were assigned to the same folds as their parent sequences, resulting in ten smaller folds nested within the Eff10k folds. This ensured that sequence-similar proteins were not shared across training, validation, and test sets.

Similarly, for the ClinVar dataset, we applied PSI-BLAST filtering to remove proteins in the test set that were sequence-similar to those in the training and validation folds, ensuring strict independence. This approach led to slightly different test subsets for our training methods ‘A’ and ‘B’ (see method description below) and ensured a strict and robust evaluation. ClinVar entries containing benign and pathogenic missense mutations with at least one star were used for additional testing. For method ‘A,’ this resulted in the CV-A set, containing 17,677 variants (15,867 benign and 1,810 pathogenic). For method ‘B,’ the CV-B set included 83,642 variants (58,159 benign and 25,483 pathogenic).

Further comparisons were conducted with AlphaMissense (AM) [9] and REVEL [8]. AlphaMissense data was retrieved from our relational PostgreSQL database [23], while REVEL data was downloaded from REVEL’s official site (https://sites.google.com/site/revelgenomics/downloads). REVEL variants were identified using Ensembl transcript IDs and GRCh38 positions, mapped with Ensembl EMBL files (version GRCh38.113, https://ftp.ensembl.org/pub/release-113/embl/homo_sapiens), utilizing custom Python scripts. Variants common to AM, REVEL, and either the CV-A or CV-B test sets were selected, resulting in the CCV-A set with 11,144 variants (1,131 benign and 10,013 pathogenic) and the CCV-B set with 46,475 variants (33,217 benign and 13,258 pathogenic).

To further assess pLM-SAV’s reliability, we examined AlphaMissense’s ambiguous set. Variants common between AM’s ambiguous category and REVEL were selected for evaluation, focusing on cases where AM struggled. Since score calculation required true labels, only ambiguous AM variants with REVEL predictions and reliable ClinVar annotations were included, forming the AMambRCV set with 5,503 variants from 2,069 proteins. This set was not independent of our training and validation sets. Variants with REVEL < 0.28 were considered likely benign, those with REVEL > 0.48 as likely pathogenic, and remaining variants were considered ambiguous. The thresholds of 0.28 and 0.48 reflect a trade-off between sensitivity and selectivity, with one typically around 75% and the other around 88%, depending on which end of the threshold is considered.

The division between ambiguous or certain predictions based on either AM or REVEL scores were performed for the independent CCV-A and CCV-B sets as well resulting in 581 ambiguous variants for CCV-A and 2,676 for CCV-B according to the AM split and 899 ambiguous variants for CCV-A and 5,404 for CCV-B according to the REVEL split.

Additionally, we compared AlphaMissense predictions using the multiplexed assays of variant effect (MAVE), also known as Deep Mutational Scanning (DMS), from the ProteinGym benchmark (version 0.1) [24]. This benchmark consists of 1.5 million variants derived from 87 assays covering 72 unique proteins.

### Training strategy

Training was conducted on the Eff10k dataset using a nested k-fold cross-validation framework, incorporating separate validation and test sets, to rigorously assess predictor performance while minimizing overfitting and data leakage. In method ‘A’, all 10 folds were utilized (k=10), whereas in method ‘B’ only folds 1 to 9 were used (k=9). Fold 0 was excluded in method ‘B’ due to its disproportionately large size, approximately seven times larger than the average of the other folds in terms of both mutations and proteins. To ensure strict independence, the test set contained no proteins sequentially similar to those in the training and validation folds to prevent data leakage.

Training was optimized using BCEWithLogitsLoss (binary cross-entropy loss combined with a sigmoid layer for numerical stability) as implemented in PyTorch [25]. To account for varying fold sizes, weighting was applied. Early stopping was implemented by monitoring the difference between training and validation losses to prevent overfitting. The optimal probability threshold for binary classification was determined by evaluating the MCC (Matthews correlation coefficient) scores across thresholds from 0.1 to 0.9 in increments of 0.01, and selecting the threshold that produced the highest respective score.

In each iteration of k-fold cross-validation, the training set was merged from k-2 folds, the validation set consisted of one fold for tuning model parameters and assessing performance, and a separate fold was held out as the test set to evaluate generalizability. This resulted in 90 models for method A and 72 for method B, with the best model selected per fold for evaluation. This nested structure ensured that every fold was used as a test set exactly once throughout the full cross-validation cycle. All reported training results represent averages across the best models, as selecting a single overall best model for a training cycle could lead to overfitting. For method ‘B’, Fold 0 combined with the corresponding test fold was also used as a hybrid prediction.

### CNN training with delta embeddings

For each protein variant, embeddings were generated for the full amino acid sequence of both the wild-type and mutant proteins using ProtT5-XL-U50 (referred to as ProtT5) and the Ankh protein language model [13,14] To capture the representational difference of a mutation, the embedding vector of the mutant sequence was subtracted from that of the wild-type sequence on a per-residue basis (Δembeddingi = embedding_WT,i_-embedding_mut,i_). Notably, while the difference in embeddings was usually most pronounced at the mutation site, significant variations were also observed in the surrounding residues within the selected window. After evaluating different window sizes (not shown), a seven-residue window comprising three residues before and three residues after the mutation was selected to best represent the localized difference between the two sequences (Figure 1). The resulting feature array for each variant consisted of 7×Δembedding, with 7×1,024 dimensions for ProtT5 and 7×1,536 for Ankh. These 7×Δembedding arrays were then used as input to a convolutional neural network (CNN) to classify each variant as neutral or effect for the training set (benign or pathogenic for ClinVar, AM and REVEL).

**Figure 1:**
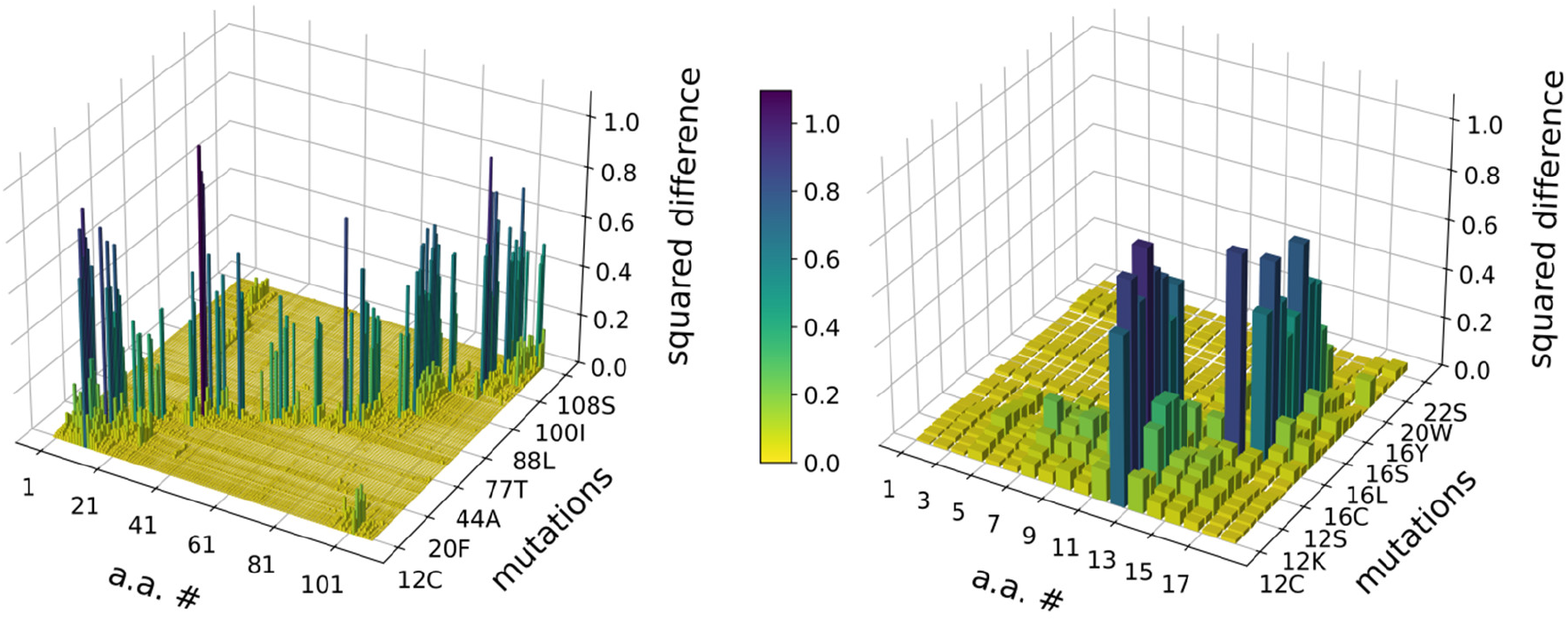
Δ-embeddings capture mutation effects beyond the mutation site. Squared differences between wild-type and mutant embeddings are shown for Cytochrome c isoform 1 (UniProt: P00044). The mutation signal is high at the mutation site but also propagates to neighboring positions, indicating local structural and functional context. These patterns highlight the suitability of convolutional neural networks (CNNs) for capturing and interpreting such localized embedding differences. (A) Squared differences across the full sequence. (B) A selected region with specific mutations is shown to enhance visibility and illustrate the propagation and decay of the signals.

Given the relatively small size of our training dataset, we opted for a compact neural network with narrow layers, offering both efficient training and fast inference times. The architecture consists of two 1D convolutional layers, each followed by a ReLU (Rectified Linear Unit) activation function, a widely used non-linearity in neural networks that outputs zero for negative values and the identity for positive values [25], scaling the output to 8 channels with a kernel size of three. After the second convolution, layer normalization is applied, followed by a residual connection, adding input to the output of a layer, to improve gradient flow, learning, and performance. The ELU (Exponential Linear Unit) activation function [25] is then used, which can produce negative outputs and helps improve learning dynamics compared to ReLU. To further reduce dimensionality, global pooling is applied – this operation condenses each feature map to a single value – before passing the output through a fully connected layer with sigmoid activation. Additionally, two dropout layers are incorporated to mitigate overfitting by randomly deactivating neurons during training. The neural network was implemented using PyTorch [25], and the class definitions are available on Zenodo (https://doi.org/10.5281/zenodo.15502498). The full pipeline and the neural network are depicted in Supplementary Figure S1.

### Performance metrics used

The performance of the predictions was assessed using standard evaluation metrics. Using the conventional definitions - TP (true positive), FP (false positive), TN (true negative), and FN (false negative) - the following metrics were calculated: (1) True positive rate (TPR), recall_effect, sensitivity = TP/(TP+FN) (2) True negative rate (TNR), recall_neutral, specificity = TN/(TN+FP) (3) Precision_effect_ = TP + TP/FP (4) Precision_neutral_ = TN/(TN+FN) (5) F1_effect_ = 2*(precision_effect_*recall_effect_)/(precision_effect_+recall_effect_) (6) F1_neutral_ = 2*(precision_neutral_ * recall_neutral_) / (precision_neutral_+recall_neutral_) (7) Accuracy = (TP+TN)/(TP+TN+FP+FN) (8) MCC = (TP*TN-FP*FN) / ((TP+FP) * (TP+FN) * (TN+FN) * (TN+FP))^-½^. The AUC (Area Under the Receiver Operating Characteristic Curve) was computed as the area under the FPR versus (1 - TNR) curve. The corrected sample standard deviation of performance was determined from the results of the best models, based on 10 models for method ‘A’ and 9 models for method ‘B’ in the case of pLM-SAV, and bootstrapping (n=1,000) was used in case of AlphaMissense and REVEL data.

For the ProteinGym v0.1 benchmark, performance was evaluated using the methodology described in the original study [24]and AlphaMissense [9]. Since this benchmark includes variants with multiple amino acid mutations, raw scores were calculated separately for each mutation within a multi-mutant variant (assuming independent effects, although this assumption may not always hold). These scores were summed to obtain a combined score. A sigmoid function was applied to the summed score, generating a single prediction per multi-mutant variant.

Spearman’s rank correlation was computed separately for each assay. The results were then averaged at the protein level, as some proteins had multiple assays. A final mean of Spearman correlation was calculated across 72 proteins in the benchmark. To ensure that a positive correlation reflects agreement between assay values and predictions, the sign of the Spearman correlation coefficients was inverted. Although some studies (including AlphaMissense) report the absolute value of Spearman correlation instead of the signed value, we followed the ProteinGym v0.1 benchmark recommendations and retained the original signed correlation. Consequently, the AlphaMissense Spearman correlation for this benchmark was recalculated for consistency with this methodology.

## Results

### Performance of delta-embedding-based predictors

We evaluated four training settings (Ankh vs. ProtT5 × method A vs. method B). The decision to handle Fold 0 separately stemmed from the observation that it was approx. 7 times larger than the other folds. The average performance metrics of the best models for each training method are presented in Figure 2 and Supplementary Table 1. In all cases, these metrics exceeded their respective statistical significance thresholds, such as 0.8 and 0.5 for AUC and MCC, respectively. Among all variations, method ‘B’ and the Ankh combination yielded the best performance. Additionally, method ‘B’ allowed for independent predictions on Eff10k Fold 0, in combination with the respective test folds. The combined predictions yielded lower scores compared to the test fold predictions, and method ‘B’ outperformed method ‘A’ overall. These observations suggest that fold 0 is inherently more difficult to predict than the other folds.

**Figure 2:**
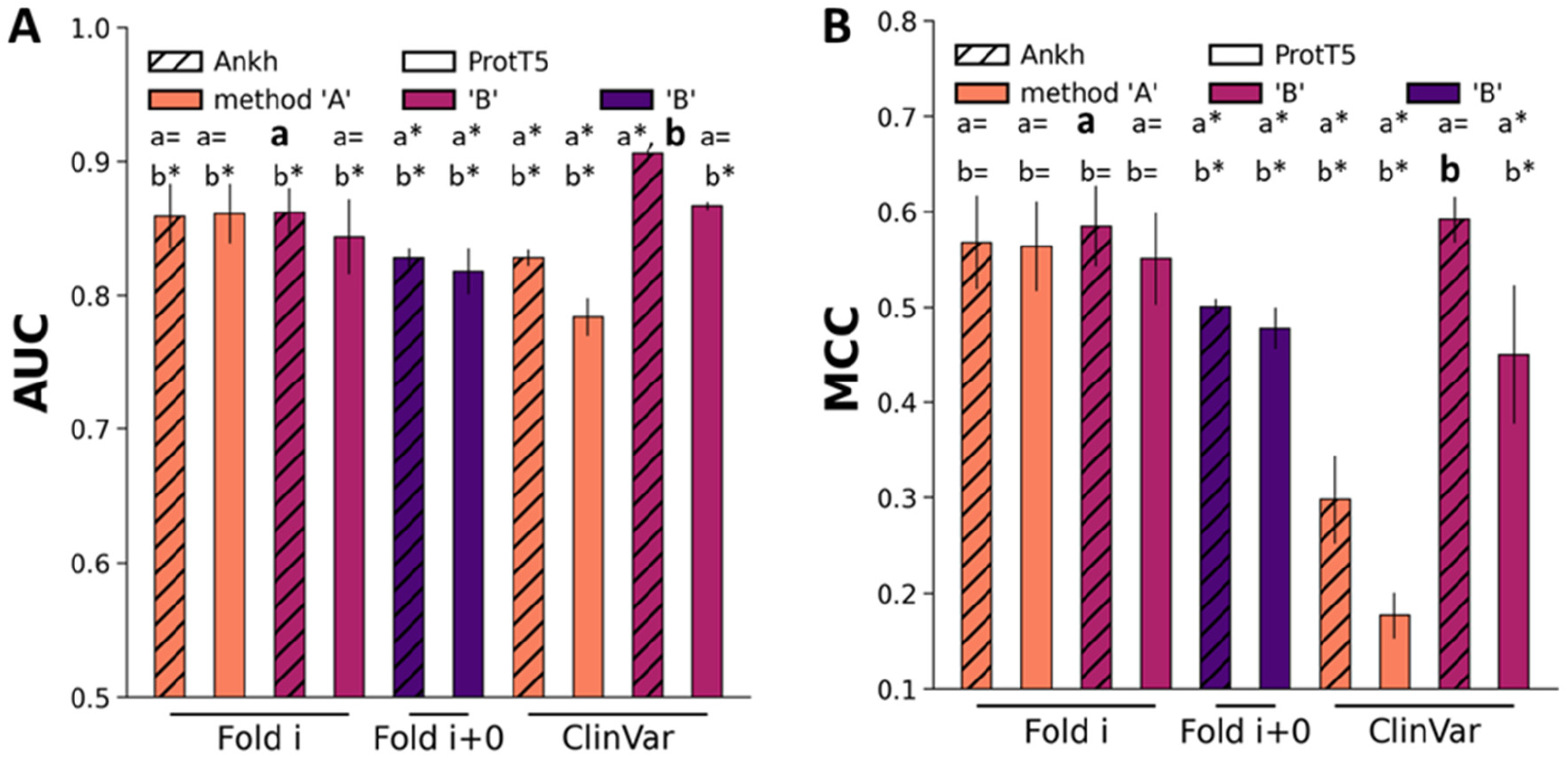
pLM-SAV performance on Eff10k sets demonstrates the effectiveness of Δ-embeddings. AUC (**A**) and MCC (**B**) values were calculated for Eff10k folds (Fold i) using method A (orange) and method B (raspberry). Additionally, evaluations for Eff10k were performed for Fold i combined with Fold 0 using method B (purple), as well as for a ClinVar-derived protein set, filtered to prevent sequence-based data leakage. The pLM-SAV evaluation was conducted using embeddings generated with Ankh (striped columns) and ProtT5 (plain columns). Values labeled with a* and b* are significantly different from values labeled with a and b as references, respectively. Differences were tested using the Wilcoxon-rank test, and were considered significant at *p* < 0.05, or non-significant (labeled with a= or b=) otherwise.

Analysis of the Eff10k dataset revealed that effect variants generally exhibit higher summed squared differences within the 7-amino acid Δ-embedding window compared to neutral variants (Supplementary Figure S2). When further separating both classes into true and false predictions by the ‘A’ and ‘B’ Ankh models, a consistent tendency emerged: correctly classified variants (true positives/negatives) align with the expected distribution of larger or smaller differences, whereas misclassified variants (false positives/negatives) often deviate from this pattern. These observations suggest that the magnitude of the wild-type versus variant embedding difference contributes to the learning process of pLM-SAV, although it is not the sole factor.

To further assess the generalization performance of our models, predictions were also made on two ClinVar datasets, CV-A (for method ‘A’) and CV-B (for method ‘B’). We created these datasets to contain only proteins that are entirely independent of the training and validation sets, without sequence similarity to any proteins in these sets. Among all tested models, only the method ‘B’/Ankh combination produced statistically significant results across all metrics in ensemble predictions (Figure 2 and Supplementary Table S1). In every case, the Ankh model outperformed ProtT5, confirming its superior ability to capture information rich sequence representations. Importantly, the superior performance of method ‘B’ suggests that Eff10k Fold 0 negatively impacts not only predictions but also the training process.

### Comparison of pLM-SAV to SAV effect predictors

To evaluate the performance of our approach, we compared it to supervised and rule-based methods. We note that our evaluation setting mirrors that of SNAP2 and VESPA, as we obtained the folds from the authors [7,10], allowing direct comparison. These methods were assessed on the PMD4k set, which is a subset of Eff10k. Therefore, we followed a careful procedure to ensure that predictions were only made for PMD4k proteins that appeared in the independent test fold, mirroring the methodology used in previous studies. No single method consistently outperformed all others across every metric (Figure 3A and Supplementary Table S2). SNAP2 achieved the highest MCC score of 0.280, while the pLM-SAV Ankh models performed nearly as well, with scores of 0.274 and 0.271, also comparable to VESPA and its variant, VESPAl. For MCC, the supervised models (pLM-SAV and VESPA) outperformed the rule-based methods. In contrast, for accuracy, the rule-based ConSeq-19-equal method, which categorizes all non-native variants at conserved positions as having an effect, achieved the highest score of 0.715 (Supplementary Table S2). These results demonstrate that Δ-embeddings alone can successfully capture the effect of single amino acid variants, producing predictions that are competitive with complex models incorporating explicit evolutionary and structural features.

**Figure 3:**
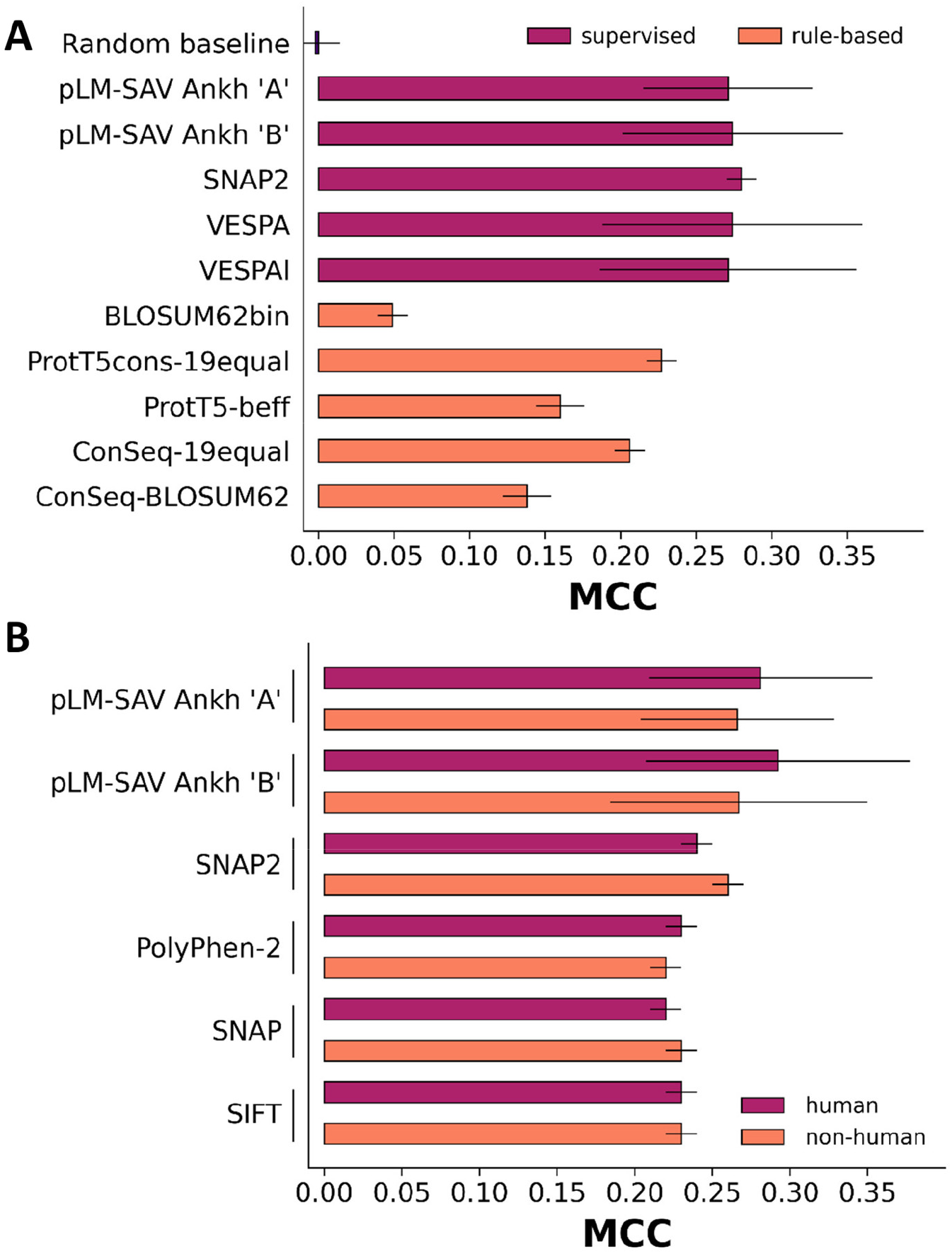
pLM-SAV performance is comparable to supervised and rule-based methods. Mean MCC values and standard errors for PMD4k (**A**) and human versus non-human split of PMD4k (**B**) datasets were collected from published sources, as detailed in the Methods and main text. pLM-SAV Ankh ‘A’ and ‘B’ were not statistically different (Wilcoxon rank-sum test, p<=0.05).

To further assess performance, we evaluated predictions on PMD4k split into human and non-human proteins, allowing a direct comparison to SNAP2’s development study [7]. This also allowed additional comparisons made with the widely used methods PolyPhen-2 [26] and SIFT [27]. As before, predictions were made only for independent test fold proteins, ensuring a fair evaluation. For human variants, the pLM-SAV ‘B’ Ankh model achieved the best scores in both MCC (0.292) and F1_neutral_ (0.497) (Figure 3B and Supplementary Table S3). For non-human variants, pLM-SAV models consistently achieved the highest scores. However, as with previous comparisons, no single method emerged as the best across all metrics.

Importantly, the mean values and standard deviations for methods other than pLM-SAV in case of PMD4k were obtained from related publications [7,10] thus statistical significance between these methods and ours could not be evaluated.

### pLM-SAV improves prediction of ambiguous AlphaMissense and REVEL variants

The performance of our pLM-SAV predictor was compared to the state-of-the-art SAV predictor AlphaMissense and to the ensemble-based REVEL method. To ensure a fair comparison, independent test sets were selected from ClinVar, including only proteins with no sequence similarity to those used in training or validation (sets CCV-A and CCV-B). These sets contained variants for which both AlphaMissense and REVEL predictions were available. Prediction scores (AUC and MCC) across the full datasets were generally lower for pLM-SAV models compared to AlphaMissense and REVEL. When the datasets were divided into ambiguous and certain prediction subsets, pLM-SAV models consistently outperformed both AlphaMissense and REVEL on the ambiguous subsets across all evaluated metrics. Using pLM-SAV method ‘A’, both Ankh and ProtT5 embeddings yielded higher AUC, MCC, and F1_effect_ scores compared to AlphaMissense on its ambiguous subset (Supplementary Table S4). In the case of the ambiguous REVEL subset, only pLM-SAV Ankh exceeded REVEL across all three metrics (AUC, MCC, F1_effect_), whereas pLM-SAV ProtT5 did not (Supplementary Table S4). Similar patterns were observed with method ‘B’, where both pLM-SAV Ankh and ProtT5 outperformed AlphaMissense on its ambiguous variants (Figure 4). For ambiguous REVEL variants under method ‘B’, pLM-SAV Ankh again outperformed REVEL in all metrics, while ProtT5 showed improvement in AUC only.

**Figure 4:**
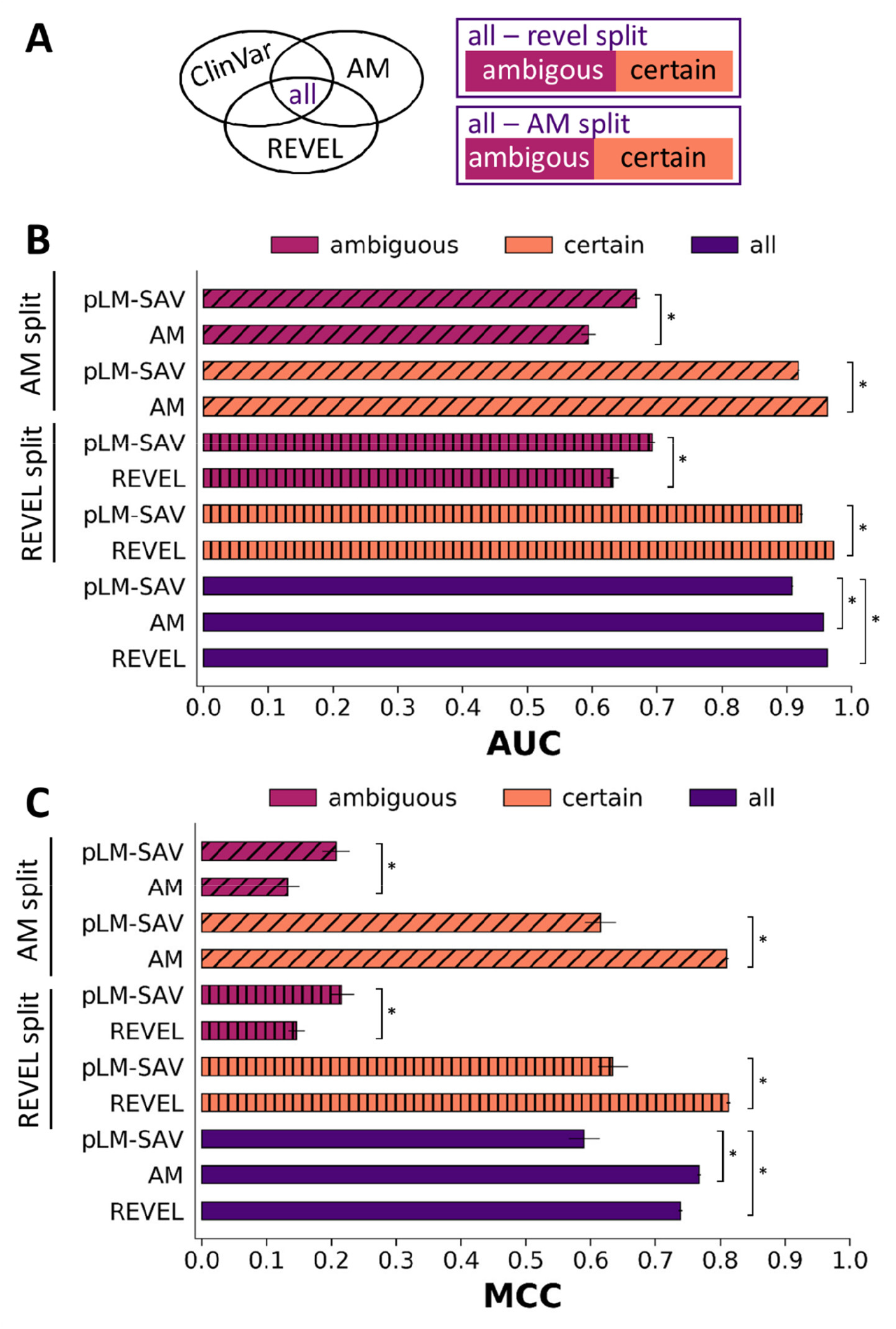
Comparison of the pLM-SAV predictions to AlphaMissense (AM) and REVEL. The test set contained proteins sequentially not similar to those in the training and validation of the pLM-SAV Ankh model (method ‘B’, set CCV-B). Metrics were calculated for ambiguous, certain and all predictions, splitting the data set according to the ambiguous/certain criteria of AlphaMissense or REVEL. Differences were tested using the Wilcoxon rank-sum test, and were considered significant at *p* < 0.05, indicated by *.

Encouraged by these promising results on the ambiguous subsets of both AlphaMissense and REVEL within the ClinVar test sets, we extended our analysis to a larger ambiguous AlphaMissense dataset (AMambRCV). To assess whether pLM-SAV could outperform these methods on a broader scale, we selected the largest possible variant set that included ambiguous AlphaMissense and REVEL predictions, labeled by ClinVar data. pLM-SAV consistently outperformed AlphaMissense, except in the case of F1_neutral_, while REVEL achieved the highest overall performance among the three methods (Figure 5 and Supplementary Table S5). Given this, for variants with ambiguous AlphaMissense predictions, it may be beneficial to refer to REVEL predictions when available.

**Figure 5:**
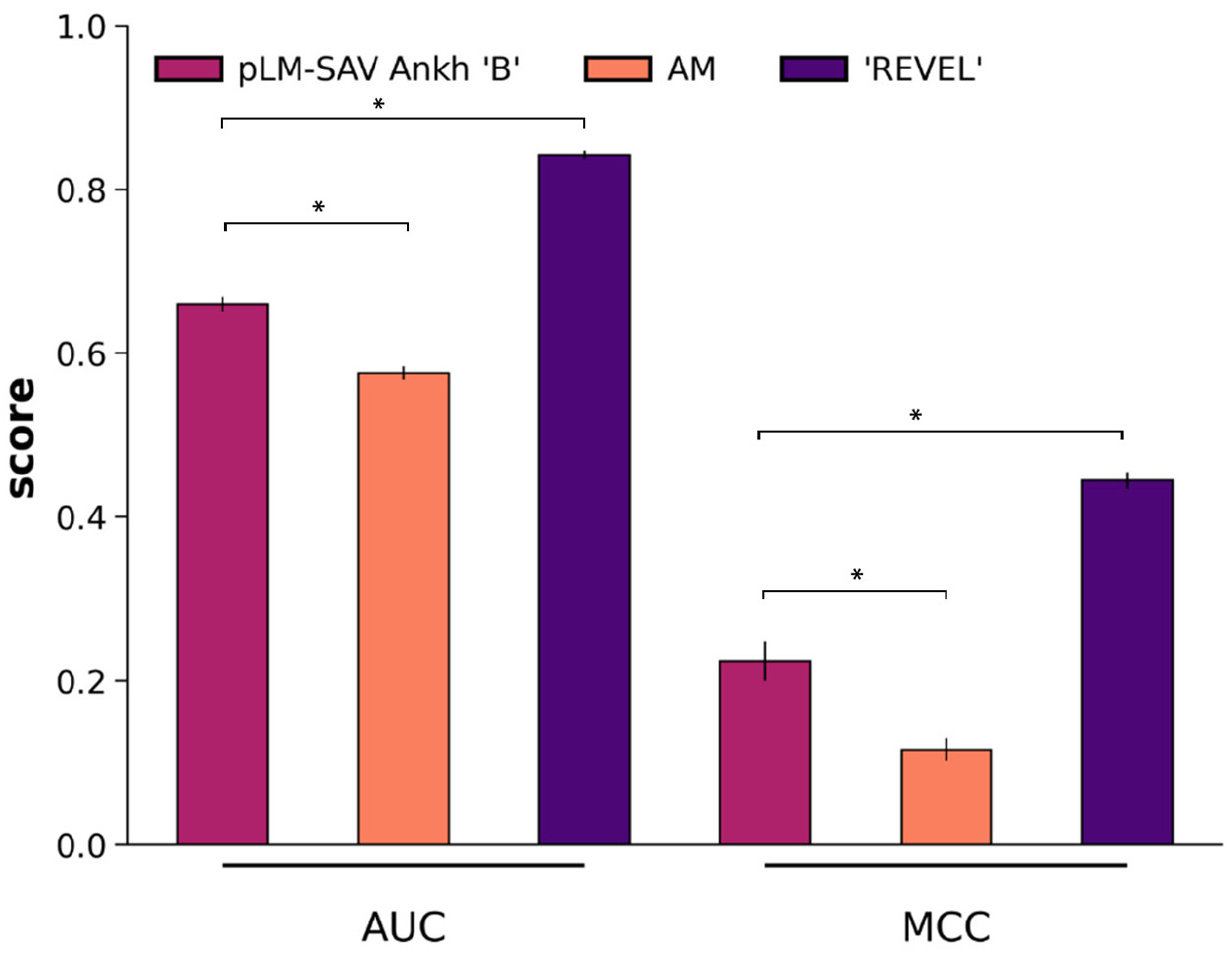
AUC and MCC scores of pLM-SAV, AlphaMissense, and REVEL predictions for the AMambRCV set. This set includes ambiguous AlphaMissense and REVEL predictions with ClinVar labels. Differences were tested using the Wilcoxon rank-sum test and considered significant at *p* < 0.05 (indicated by *).

**Figure 5:**
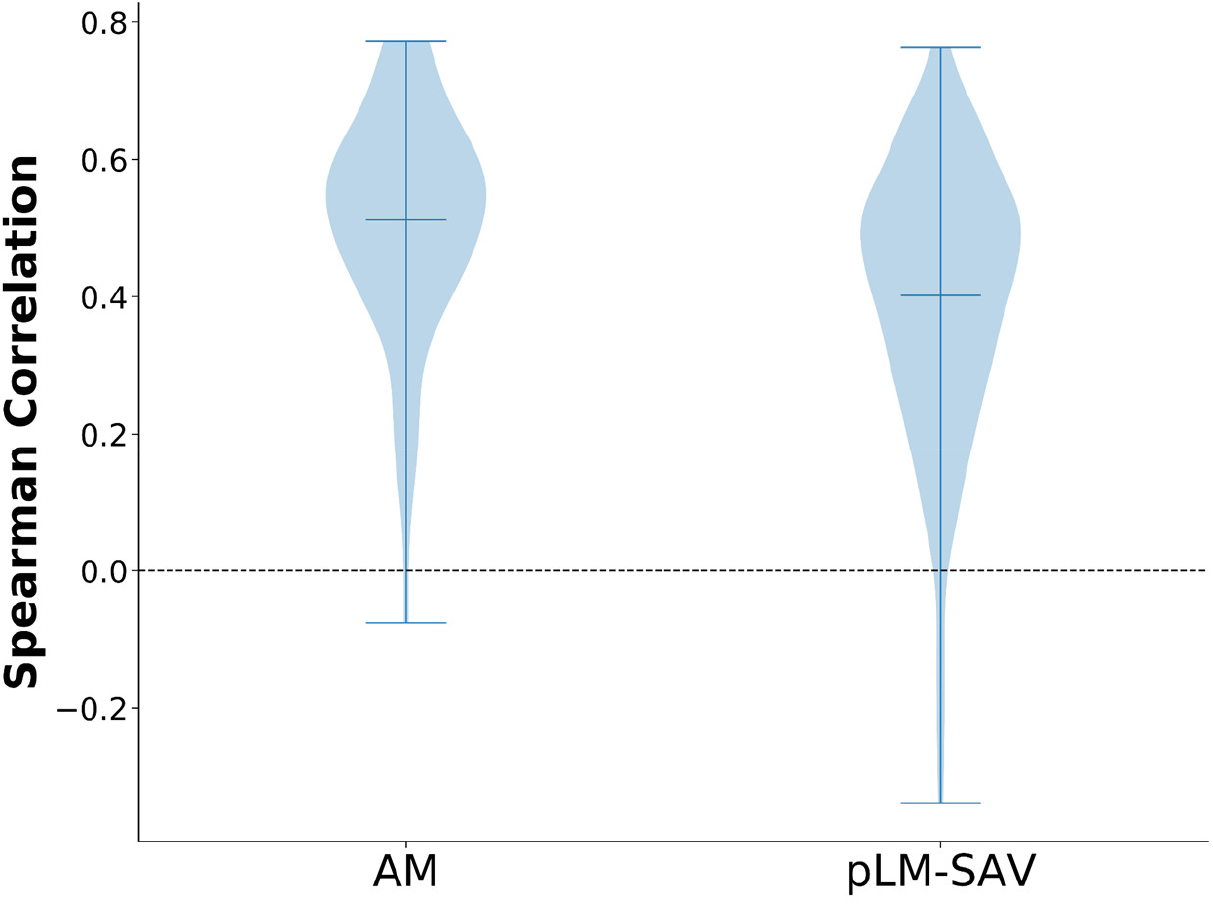
Comparison of the Spearman’s correlation for AlphaMissense and pLM-SAV (method ‘B’ with Ankh).

### Comparison with the ProteinGym v0.1 benchmark

To evaluate the predictive power of pLM-SAV on a MAVE (or DMS) benchmark, where mutation effects are represented by continuous values rather than binary labels, we used ProteinGym version 0.1 [NO_PRINTED_FORM], which was also employed in the AlphaMissense study. Although the best-performing pLM-SAV model yielded a lower average Spearman rank correlation (0.382) compared to AlphaMissense (0.512), pLM-SAV performed in a similar range as the unsupervised ESM-1b on this benchmark (0.358) [28].

## Discussion

In this study, we demonstrated that a simple Δ-embedding-based architecture, pLM-SAV, can achieve performance comparable to more complex SAV effect prediction methods. By leveraging the representational power of protein language models (pLMs), and focusing on the difference between wild-type and mutant embeddings, we showed that even a lightweight convolutional neural network can extract informative features for binary classification of SAVs.

Our results confirm the viability of using Δ-embeddings for SAV effect prediction. The best-performing pLM-SAV models (particularly those using the Ankh language model and trained with method ‘B’) consistently achieved strong predictive performance on the Eff10k test folds and on independent ClinVar subsets (Figure 2). Notably, pLM-SAV showed solid performance across both human and non-human proteins in the PMD4k benchmark and performed in a similar range as VESPA and SNAP2 (Figure 3). This suggests that Δ-embeddings alone can efficiently capture the functional effects of mutations, even without explicit structural, evolutionary, or annotation-based features. This distinguishes pLM-SAV from approaches like VESPA, which combine pLM embeddings with substitution matrices and manually curated features such as BLOSUM scores and amino acid probabilities. In contrast, pLM-SAV relies solely on Δ-embeddings, enabling a leaner and more interpretable architecture, while also achieving comparable or shorter run times (Supplementary Table S6). It is important to note that while we highlight the strong performance of pLM-SAV with its lightweight setup, developing and training protein language models is a complex and computationally expensive task. These models encode information with a level of complexity potentially comparable to that extracted from MSAs and other biological data sources.

While pLM-SAV did not outperform top-tier methods such as AlphaMissense [9]or REVEL [8] across the full dataset evaluations (Figure 4), it showed a distinct advantage in predicting the effects of variants within ambiguous regions (Figure 4-5), where AlphaMissense and REVEL provide low-confidence or uncertain outputs. In these difficult subsets, pLM-SAV outperformed both AlphaMissense and REVEL in most metrics, especially when using Ankh embeddings. This suggests that Δ-embeddings provide a complementary signal that can resolve uncertain cases, and therefore could be integrated into ensemble models or meta-predictors to improve overall reliability. We leveraged this insight and evaluated how pLM-SAV and REVEL perform on AlphaMissense predictions with ambiguous scores (Figure 5). Since REVEL outperformed both pLM-SAV and AlphaMissense on this subset, we incorporated REVEL predictions into our web application (alphamissense.hegelab.org) [23] to enhance its utility.

It is important to emphasize that our approach is trained on labeled data derived from the Eff10k dataset [7], where class labels indicate whether a mutation has any measurable effect, regardless of its pathogenicity. In contrast, the ClinVar-based evaluation relies on a strict benign versus pathogenic distinction. This mismatch in classification categories could partly explain the lower performance observed in some ClinVar test sets, especially in the CV-A subset, which is significantly smaller and more label-imbalanced than CV-B. Lastly, our model’s performance on the ProteinGym benchmark further supports the relevance of Δ-embeddings. Although pLM-SAV did not reach the performance of AlphaMissense in this benchmark, it matched the unsupervised ESM-1b model [28], suggesting that it captures biologically meaningful variation despite being trained on binary labels. These observations highlight that current benchmarks may impose a performance ceiling due to dataset limitations such as label ambiguity, imbalance, and inconsistent annotation standards. However, we believe that further improvements remain possible through better handling of ambiguous variants and integration of complementary features such as structure-aware representations.

While pLM-SAV was developed for single amino acid variants, the approach could be extended to more complex variants such as insertions, deletions, and fusions. In these cases, embedding pairs cannot be subtracted within the altered region, but Δ-embeddings from surrounding residues could still be computed and aggregated for classification. Given the contextual encoding of pLMs, effects of such variants are likely to manifest outside the mutated region as well. However, using larger windows may introduce noise, and the lack of high-quality training data currently limits practical implementation.

Interestingly, the Δ-embedding signal often extends beyond the mutated residue (Figure 1), suggesting that pLM-based representations capture contextual dependencies. This raises the possibility that predictions may be modulated by other sequence variations – benign or otherwise – appearing elsewhere in the protein. However, the usually low Δ-embedding values at positions distant from the mutation site (Figure 1), together with our preliminary results (Supplementary Table S7), suggest that such distant residues do not systematically influence the model’s output.

Overall, pLM-SAV offers a fast and easily deployable solution for SAV effect prediction. It generalizes well across datasets, performs robustly on ambiguous variants, and requires fewer input features than traditional methods. We expect that future developments incorporating delta-embeddings into larger architectures, or combining them with structural or evolutionary signals, may further improve predictive accuracy and clinical relevance.

## Data and code availability

The data and code used in this article can be downloaded from Zenodo, https://doi.org/10.5281/zenodo.15502498. The associated web server is freely available at https://alphamissense.hegelab.org.

## Acknowledgements

Thanks to Burkhard Rost (Technical University Munich) for the Eff10k dataset and his colleague, Céline Marquet for discussions.

## CRediT authorship contribution statement

O. Gereben: Writing – original draft, Investigation, Methodology; H. Tordai: Data curation, Visualization, Writing – review and editing; L. Khamisi: Data curation, Writing – review and editing; E. Qorri: Methodology, Writing – review and editing; T. Hegedűs: Conceptualization, Investigation, Methodology, Funding acquisition, Writing – original draft.

## Declaration of generative AI and AI-assisted technologies in the writing process

During the preparation of this work the authors used ChatGPT in order to improve language. After using this tool/service, the authors reviewed and edited the content as needed and take full responsibility for the content of the publication.

