## Supplemental Figures and Tables for "pLM-SAV: A Δ-Embedding Approach for Predicting Pathogenic Single Amino Acid Variants"

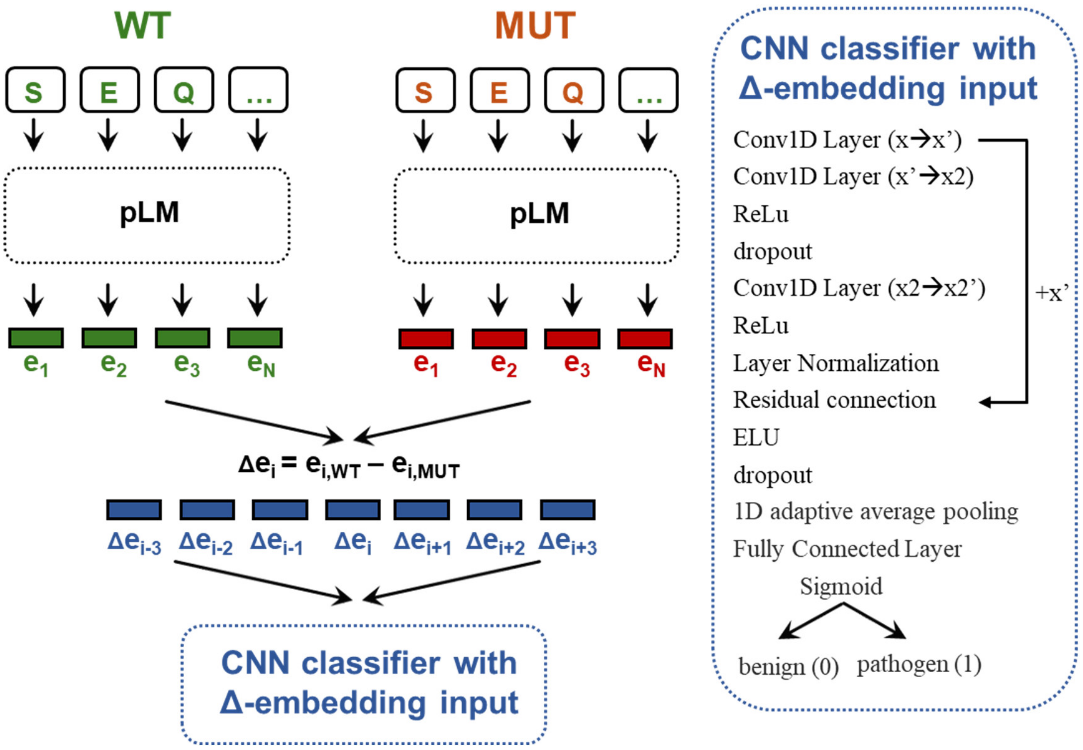


**Figure S1: Schematic representation of pLM-SAV architecture.** WT: wild type, MUT: mutant, pLM: protein language model, $e_i$: embedding of a residue, Δe: delta embedding calculated as $embedding_{i,WT}$ - $embedding_{i,MUT}$, CNN: convolutional neural network, ReLu: rectified linear unit, ELU: exponential linear unit. Please see detailed description in the main text.

**Table S1: pLM-SAV performance on Eff10k sets demonstrates the effectiveness of Δ-embeddings.** AUC and MCC values were calculated for Eff10k folds (Fold i) using method A and method B. Additionally, evaluations for SNAP2 were performed for Fold i combined with Fold 0 using method B, as well as for a ClinVar-derived protein set, filtered to prevent sequence-based data leakage. The pLM-SAV evaluation was conducted using embeddings generated with Ankh and ProtT5. Mean values with standard deviation are shown. Values labeled with a* and b* are significantly different from values labeled with a and b as references, respectively. Differences were tested using the Wilcoxon-rank test, and were considered significant at $p < 0.05$, or non-significant (labeled with a= or b=) otherwise.

| Method | AUC | MCC |
|---|---|---|
| **'A' Ankh Fold i** | 0.859 ± 0.024 (a=, b*) | 0.568 ± 0.049 (a=, b=) |
| **'A' ProtT5 Fold i** | 0.861 ± 0.022 (a=, b*) | 0.564 ± 0.047 (a=, b=) |
| **'B' Ankh Fold i** | 0.862 ± 0.018 (a, b*) | 0.585 ± 0.042 (a, b=) |
| **'B' ProtT5 Fold i** | 0.844 ± 0.028 (a=, b*) | 0.551 ± 0.048 (a=, b=) |
| **'B' Ankh Fold i + 0** | 0.828 ± 0.007 (a*, b*) | 0.501 ± 0.008 (a*, b*) |
| **'B' ProtT5 Fold i + 0** | 0.818 ± 0.017 (a*, b*) | 0.478 ± 0.022 (a*, b*) |
| **'A' Ankh ClinVar** | 0.828 ± 0.006 (a*, b*) | 0.298 ± 0.046 (a*, b*) |
| **'A' ProtT5 ClinVar** | 0.784 ± 0.014 (a*, b*) | 0.177 ± 0.024 (a*, b*) |
| **'B' Ankh ClinVar** | 0.906 ± 0.003 (a*, b) | 0.592 ± 0.024 (a=, b) |
| **'B' ProtT5 ClinVar** | 0.867 ± 0.003 (a=, b*) | 0.450 ± 0.073 (a*, b*) |

**Table S2 pLM-SAV performance is comparable to supervised methods. (A)** Mean MCC and accuracy values and standard errors for PMD4k were collected from published sources, as detailed in the Methods and main text. pLM-SAV Ankh 'A' and 'B' were not statistically different (Wilcoxon rank-sum test, p<=0.05). Mean values with standard deviation are shown. Difference compared to set 'a' was tested using the Wilcoxon-rank test, and was considered non-significant at $p < 0.05$ labeled with a=).

| Method | MCC | Accuracy |
|---|---|---|
| **Random baseline** | -0.002 ± 0.611 | 0.611 ± 0.41 |
| **Supervised** | | |
| **pLM-SAV Ankh 'A'** | 0.271 ± 0.696[(a=)] | 0.696 ± 0.038[(a=)] |
| **pLM-SAV Ankh 'B'** | 0.274 ± 0.694[(a)] | 0.694 ± 0.036[(a)] |
| **SNAP2** | 0.280 ± 0.707 | 0.707 ± 0.39 |
| **VESPA** | 0.274 ± 0.635 | 0.635 ± 0.43 |
| **VESPAl** | 0.271 ± 0.63 | 0.630 ± 0.43 |
| **Rule-based** | | |
| **BLOSUM62bin** | 0.049 ± 0.562 | 0.562 ± 0.43 |
| **ProtT5cons-19equal** | 0.227 ± 0.686 | 0.686 ± 0.41 |
| **ProtT5-beff** | 0.16 ± 0.523 | 0.523 ± 0.43 |
| **ConSeq-19equal** | 0.206 ± 0.715 | **0.715 ± 0.39** |
| **ConSeq BLOSUM62** | 0.138 ± 0.543 | 0.543 ± 0.43 |

**Table S3 pLM-SAV performance is comparable to rule-based methods.** Mean MCC, $F1_{effect}$, $F1_{neutral}$ and accuracy for the human versus non-human split of PMD4k datasets were collected from published sources, as detailed in the Methods and main text. pLM-SAV Ankh 'A' and 'B' were not statistically different (Wilcoxon rank-sum test, p<=0.05) neither in the human (a) or in the non-human (b) split. Mean values with standard deviation are shown.

| Method | MCC | $F1_{effect}$ | $F1_{neutral}$ | Accuracy |
|---|---|---|---|---|
| | | **human** | | |
| **pLM-SAV Ankh 'A'** | 0.281 ± 0.072[(a=)] | 0.766 ± 0.038[(a=)] | 0.492 ± 0.053[(a=)] | 0.681 ± 0.043[(a=)] |
| **pLM-SAV Ankh 'B'** | **0.292 ± 0.085**[(a)] | 0.763 ± 0.037[(a)] | **0.497 ± 0.058**[(a)] | 0.678 ± 0.045[(a)] |
| **SNAP2** | 0.24 ± 0.01 | 0.780 ± 0.6 | 0.463 ± 1.3 | 0.688 ± 0.7 |
| **PolyPhen-2** | 0.23 ± 0.01 | **0.784 ± 0.4** | 0.451 ± 1.1 | **0.689 ± 0.5** |
| **SNAP** | 0.22 ± 0.01 | 0.749 ± 0.5 | 0.467 ± 1.1 | 0.658 ± 0.6 |
| **SIFT** | 0.23 ± 0.01 | 0.722 ± 0.6 | 0.490 ± 1.0 | 0.636 ± 0.6 |
| | | **non-human** | | |
| **pLM-SAV Ankh 'A'** | 0.266 ± 0.062[(b=)] | 0.789 ± 0.036[(b=)] | 0.459 ± 0.065[(b=)] | 0.699 ± 0.040[(b=)] |
| **pLM-SAV Ankh 'B'** | **0.267 ± 0.083**[(b)] | 0.787 ± 0.033[(b)] | **0.461 ± 0.077**[(b)] | 0.697 ± 0.039[(b)] |
| **SNAP2** | 0.26 ± 0.01 | 0.799 ± 0.3 | 0.458 ± 0.8 | 0.707 ± 0.4 |
| **PolyPhen-2** | 0.22 ± 0.01 | 0.771 ± 0.4 | 0.447 ± 0.8 | 0.676 ± 0.5 |
| **SNAP** | 0.23 ± 0.01 | 0.772 ± 0.3 | 0.455 ± 0.9 | 0.679 ± 0.5 |
| **SIFT** | 0.23 ± 0.01 | 0.770 ± 0.3 | 0.458 ± 0.8 | 0.677 ± 0.4 |

**Table S4 Comparison of the pLM-SAV predictions to AlphaMissense (AM) and REVEL.** The test set contained proteins sequentially not similar to those in the training and validation of the pLM-SAV Ankh model (method 'A' and 'B'). Metrics were calculated for certain, ambiguous, and all predictions, splitting the data set according to the ambiguous/certain criteria of AlphaMissense and REVEL. Values labeled with a*…j* are significantly different from values labeled with a…j as references, respectively. Differences were tested using the Wilcoxon-rank test, and were considered significant at $p < 0.05$.

| method | AUC | MCC | F1effect |
|---|---|---|---|
| 'A' Ankh AM ambiguous | 0.651 ± 0.009(a) | 0.147 ± 0.019(a) | 0.258 ± 0.011(a) |
| 'A' ProtT5 AM ambiguous | 0.558 ± 0.013(a*) | 0.048 ± 0.022(a*) | 0.215 ± 0.007(a*) |
| AM ambiguous | 0.531 ± 0.038(a*) | 0.005 ± 0.042(a*) | 0.181 ± 0.031(a*) |
| 'A' Ankh AM certain | 0.857 ± 0.003(b) | 0.327 ± 0.051(b) | 0.364 ± 0.051(b) |
| 'A' ProtT5 AM certain | 0.822 ± 0.011(b*) | 0.199 ± 0.025(b*) | 0.256 ± 0.018(b*) |
| AM certain | 0.957 ± 0.003(b*) | 0.687 ± 0.011(b*) | 0.711 ± 0.010(b*) |
| 'A' Ankh REVEL ambiguous | 0.677 ± 0.006(c) | 0.191 ± 0.036(c) | 0.431 ± 0.017(c) |
| 'A' ProtT5 REVEL ambiguous | 0.611 ± 0.007(c*) | 0.083 ± 0.020(c*) | 0.386 ± 0.007(c*) |
| REVEL ambiguous | 0.635 ± 0.020(c*) | 0.157 ± 0.032(c*) | 0.398 ± 0.024(c*) |
| 'A' Ankh REVEL certain | 0.865 ± 0.003(d) | 0.320 ± 0.049(d) | 0.343 ± 0.051(d) |
| 'A' ProtT5 REVEL certain | 0.836 ± 0.011(d*) | 0.198 ± 0.025(d*) | 0.237 ± 0.018(d*) |
| REVEL certain | 0.961 ± 0.003(d*) | 0.741 ± 0.011(d*) | 0.763 ± 0.010(d*) |
| 'A' Ankh all | 0.848 ± 0.003(e) | 0.315 ± 0.047(e) | 0.355 ± 0.047(e) |
| 'A' ProtT5 all | 0.809 ± 0.011(e*) | 0.192 ± 0.024(e*) | 0.253 ± 0.017(e*) |
| AM all | 0.950 ± 0.003(e*) | 0.628 ± 0.010(e*) | 0.657 ± 0.010(e*) |
| REVEL all | 0.950 ± 0.003(e*) | 0.648 ± 0.011(e*) | 0.682 ± 0.010(e*) |
| 'B' Ankh AM ambiguous | 0.668 ± 0.006(f) | 0.207 ± 0.021(f) | 0.506 ± 0.009(f) |
| 'B' ProtT5 AM ambiguous | 0.631 ± 0.006(f*) | 0.128 ± 0.038(f*) | 0.471 ± 0.003(f*) |
| AM ambiguous | 0.594 ± 0.011(f*) | 0.132 ± 0.019(f*) | 0.441 ± 0.013(f*) |
| 'B' Ankh AM certain | 0.918 ± 0.002(g) | 0.615 ± 0.024(g) | 0.729 ± 0.017(g) |
| 'B' ProtT5 AM certain | 0.885 ± 0.004(g*) | 0.473 ± 0.078(g*) | 0.634 ± 0.053(g*) |
| AM certain | 0.963 ± 0.001(g*) | 0.810 ± 0.003(g*) | 0.865 ± 0.002(g*) |
| 'B' Ankh REVEL ambiguous | 0.692 ± 0.005(h) | 0.216 ± 0.019(h) | 0.376 ± 0.013(h) |
| 'B' ProtT5 REVEL ambiguous | 0.643 ± 0.005(h*) | 0.135 ± 0.022(h*) | 0.326 ± 0.009(h*) |
| REVEL ambiguous | 0.632 ± 0.009(h*) | 0.146 ± 0.013(h*) | 0.335 ± 0.010(h*) |
| 'B' Ankh REVEL certain | 0.923 ± 0.002(i) | 0.634 ± 0.023(i) | 0.749 ± 0.016(i) |
| 'B' ProtT5 REVEL certain | 0.890 ± 0.004(i*) | 0.490 ± 0.079(i*) | 0.656 ± 0.052(i*) |
| REVEL certain | 0.972 ± 0.001(i*) | 0.813 ± 0.003(i*) | 0.869 ± 0.002(i*) |
| 'B' Ankh all | 0.908 ± 0.002(j) | 0.590 ± 0.024(j) | 0.713 ± 0.016(j) |
| 'B' ProtT5 all | 0.873 ± 0.004(j*) | 0.454 ± 0.075(j*) | 0.624 ± 0.050(j*) |
| AM all | 0.957 ± 0.001(j*) | 0.767 ± 0.003(j*) | 0.835 ± 0.002(j*) |
| REVEL all | 0.963 ± 0.001(j*) | 0.739 ± 0.003(j*) | 0.814 ± 0.002(j*) |

**Table S5 AUC and MCC scores of pLM-SAV, AlphaMissense, and REVEL predictions for the largest possible variant set that included ambiguous AlphaMissense and REVEL predictions.** Differences compared to set 'a' were tested using the Wilcoxon rank-sum test and considered significant at $p < 0.05$ (indicated by a*).

| method | AUC | MCC |
|---|---|---|
| **'B' Ankh** | 0.660 ± 0.009[(a)] | 0.224 ± 0.024[(a)] |
| **AM** | 0.576 ± 0.008[(a*)] | 0.116 ± 0.014[(a*)] |
| **REVEL** | 0.842 ± 0.005[(a*)] | 0.445 ± 0.010[(a*)] |

**Table S6 Effect of secondary mutations on pLM-SAV predictions.** We evaluated whether additional mutations at secondary sites influence the predicted pathogenicity of a primary mutation. For CFTR (UniProt: P13569), the ΔF508 deletion was approximated by the F508G substitution, as pLM-SAV does not support deletions. The F508G mutation was predicted to be pathogenic (probability: 0.800) and this prediction remained unchanged even in the presence of two different sets of known rescue mutations (3PT and 6SS without ΔRI, Soya *et al.* Nat Commun 14, 6868, 2023). For human LILRB4 (UniProt: Q8NHJ6), we examined the mutation of Cys144 – part of a disulfide bridge in the Ig-like domain – which was predicted to be pathogenic. We hypothesized that a compensatory mutation at the corresponding cysteine (C195) might rescue the prediction. Several combinations were tested, including substitutions to charged residues (D) and small residues (Ala, Gly), but none resulted in a reduction in predicted pathogenicity. All these results suggest that such secondary mutations do not systematically influence the model's output.

| protein | mutation | secondary site mutation(s) | probability |
|---|---|---|---|
| CFTR | F508G | none | 0.800 |
| | F508G | S492P, A534P, I539T | 0.803 |
| | F508G | M470V, S492P, S495P, A534P, I539T, R555K | 0.803 |
| LILRB4 | C144K | none | 0.913 |
| | C144K | C195D | 0.908 |
| | C144A | none | 0.904 |
| | C144A | C195A | 0.879 |
| | C144G | none | 0.904 |
| | C144G | C195A | 0.897 |

**Table S7 pLM-SAV prediction speed compared to VESPAl.** The only available benchmark data for VESPA was for its lightweight version, VESPAl. Under this comparison, the pLM-SAV ‘A’ models achieve similar prediction speeds to VESPAl, while the ‘B’ models are more than twice as fast. The pLM-SAV speed tests were performed on an NVIDIA GeForce RTX 2080 Ti, whereas the VESPAl benchmark was run on an NVIDIA Quadro RTX 8000.

| method | N(models) | N(variants) | N(predictions) | time, s | prediction/s |
|---|---|---|---|---|---|
| VESPAl | 10 | 380,000 | 3,800,000 | 2,400 | 1,583 |
| ‘A’ Ankh | 10 | 17,677 | 176,770 | 130 | 1,360 |
| ‘A’ protT5 | 10 | 17,677 | 176,770 | 125 | 1,414 |
| ‘B’ Ankh | 9 | 83,642 | 752,778 | 180 | 4,182 |
| ‘B’ protT5 | 9 | 83,642 | 752,778 | 165 | 4,562 |
